# The NTU-DSI-122 template as a flexible platform for fiber orientation distribution registration of diffusion microstructure into stereotaxic space

**DOI:** 10.1101/2025.03.10.642453

**Authors:** Benjamin T. Newman, John D. Van Horn, T. Jason Druzgal

## Abstract

The registration of neuroimaging subjects to a common stereotaxic space allows for comparisons between timepoints, subjects, or with reference atlases of regions of interest. Typically, this has been performed by computing a similarity metric between the voxel-wise intensity values in the subject image and a reference image. Diffusion MRI is a method that provides far more detailed voxel-wise information than a simple scalar intensity value. This allows for the potential to use this additional information to perform registration with added within-tissue contrast. In this study, we present a novel use of the NTU-DSI-122 template as a fiber orientation distribution (FOD) template, for the purpose of registering subject dMRI images to stereotaxic space. The reliability and accuracy of this FOD-based registration method are compared to the intensity-based registration method ANTs by registering cellular microstructure maps from two separate cohorts to the NTU-DSI-122 template. The stochastic FOD-based method significantly outperformed the stochastic intensity-based metric on reliability and was able to more consistently register the same subject multiple times independently. The FOD-based method also significantly outperformed the intensity-based metric on registration accuracy by more completely aligning the microstructure maps to the template as measured by the Sorenson-Dice coefficient at multiple percentile thresholds. The NTU-DSI-122 template has the additional benefit of including multiple b-value shells between a wide range of feasible acquisition schemes, making the platform a flexible option for registering acquisitions of varying quality, including clinically acquired data.

## Introduction

Registering subjects to a common template space is a necessary step in almost any neuroimaging study performing group-level comparisons. A straightforward process for doing so aligns the brains of each acquisition in a study cohort from the acquired ‘native’ space to a ‘common’ template space. This common space is intended to achieve voxel-voxel correspondence between images and allow for comparison between subjects without interference from factors such as head orientation in the scanner or naturally occurring anatomical variation.

The idea of a stereotaxic common space that could allow for images, atlases, and ROIs from different studies to be applied in a standardized manner gained attention with the definition of Talairach space-registered atlases(Collins et al., 1994), and the release of stereotaxic templates such as the Colin 27 Average Brain(Fonov et al., 2011), and the MNI 152 linear template(Holmes et al., 1998).

While manual segmentation by multiple experienced raters was once viewed as the gold standard in anatomical parcellation(Desikan et al., 2006), this procedure is not always available or practical for large volume multi-site cohorts. The Adolescent Brain Cognitive Development (ABCD) cohort, for example, contains imaging across multiple modalities and multiple timepoints for over 11,000 subjects, far too many to reasonably manually segment ROIs in a timely and cost efficient manner(Casey et al., 2018; Volkow et al., 2018). Automated methods are thus necessary to align subject brain scans collected in their native space with a suitable reference template in order to apply an atlas in an unbiased manner.

As the brain differs in size and proportion between subjects(Allen et al., 2002; Reardon et al., 2018) registration algorithms must be able to warp the brain along multiple axes in order to achieve an accurate alignment. This is typically performed using an algorithm with 12 degrees of freedom: rotation, translation, scale, and shear in a non-linear affine transform based on mutual information(Maes et al., 1997). This process is most straightforward when dealing with T1-weighted structural images of subjects because the voxel intensity values create a consistent contrast between cortex, white matter (WM), and CSF (Mugler III & Brookeman, 1990). One of the most widely applied registration algorithms in neuroimaging, the ‘SyN’ model implemented in the program ANTs(Avants et al., 2011, 2014), utilizes a diffeomorphic regularization to minimize the intensity difference between images (by maximizing a cross-correlation metric), combined with assistance from anatomical priors(Avants et al., 2011). This registration approach is highly reproducible(Avants et al., 2011), and has performed well against other methods in open competition(Fu et al., 2017; Klein et al., 2009).

Diffusion MRI (dMRI) images can present a particular challenge for this process, as each individual gradient direction may have different voxels with high and low intensity based on gradient angle and subject positioning. The use of multiple b-value shell acquisitions provides an additional complication, as each individual voxel may have different intensity at each shell in addition to global intensity shifts dependent on the cellular tissue contents within each voxel(Jeurissen et al., 2014). One solution to this problem has been to register the b=0 s/mm^2^ volumes collected with the diffusion images to a T2-weighted image due to the similar anatomical contrast present in both images(Huang et al., 2008; McLaughlin et al., 2007; Pierpaoli et al., 2010). However this method neglects the strength of dMRI’s ability to provide detailed information *within* tissue compartments, especially WM and areas where myelinated axons extend into the cortex.

The voxel-wise image intensity between two WM fiber bundles may not be greatly different at b=0 (i.e., they *lack* within tissue contrast), thus intensity-based algorithms may have lower sensitivity toward within tissue location. However the *orientation* of those WM fiber bundles can vary, potentially including multiple directions within a single voxel depending on analysis method(Jeurissen et al., 2013). The presence of these crossing fibers complicates many common DTI derived metrics used in registration, such as fractional anisotropy based-registration(Farquharson et al., 2013; Oouchi et al., 2007). Accounting for this orientation information during registration is important for tractography and reconstructing accurate connectomes. Additionally, some measures of cellular microstructure, such as free water signal fractions, have a small value range (between 0-1 in this case), making the measure sensitive to even small changes in value or introduction of noise. Symmetric diffeomorphic registration via cross-correlation of the spherical harmonic coefficients has previously been demonstrated as a means of accurately co-registering WM fiber orientation distributions (FODs) within a cohort(Raffelt et al., 2011). This method allows for crossing fiber tracts to be registered between subjects, and can create group average templates composed of information from each subject in a particular study cohort. However, many neuroimaging methods such as connectome building or automated atlas fitting require registration to an existing template specifically in stereotaxic space. An additional requirement for any FOD template to achieve effective widespread use is for the FODs to be available for calculation from multiple b-values. Different b-values have greatly different FOD amplitudes(Tournier et al., 2013) and WM fiber bundle signal varies between b-value shells(Genc et al., 2020), and constrained spherical deconvolution algorithms(Newman et al., n.d.).

These two requirements: 1) that a diffusion FOD template be located in stereotaxic space, and 2) that the template be adaptable to a variety of b-values in order to match collected data, are both met by the NTU-DSI-122 template(Hsu et al., 2015). This template was developed by combining diffusion spectrum images obtained from 122 individual subjects through a multi-step registration procedure. The result is a template fit to ICBM space with multiple b-values up to b=4000 s/mm^2^. Though designed as a diffusion spectrum template the number of b-values at each shell are suitable for extraction and FOD calculation. It is then possible to register subject FOD images or group level template images with the NTU-DSI-122 at an appropriate b-value for parcellation and analysis. This study introduces the NTU-DSI-122 as a candidate FOD template in stereotaxic space for the registration of dMRI images. The utility of this method is compared in two experiments to a commonly used standard registration method for both reliability (Experiment 1) and accuracy (Experiment 2) with a specific focus on the registration of dMRI derived measures of cellular microstructure.

## Methods

### Template

The NTU-DSI-122 was developed by combining 122 individual subjects (61 male, age 27.97 ± 5.25 years, ranging from 19-40 years-old) diffusion images through a two-step registration procedure incorporating structural and diffusion weighted registration(Hsu et al., 2015). This process involved 1) creating a mean tissue probability map from all input subjects, and aligning that mean image to stereotaxic space. 2) Aligning each subject’s diffusion spectrum image and q-space information before averaging each subject to construct the final template (Hsu et al., 2012).

### Experiment 1: Assessment of Reliability

To assess the reliability of each transform method the same identical registration was performed repeatedly for a number of images and the results were compared for consistency. We collected 5 diffusion MRIs from healthy controls (all male) using a Siemens Prisma 3T scanner with an isotropic voxel size of 1.7⨉1.7⨉1.7 mm, TE=70 ms and TR=2900 ms; 10 b=0 images and 64 gradient directions at both b=1500 s/mm^2^ and b=3000 s/mm^2^. Each image set was analyzed using MSMT-CSD(Jeurissen et al., 2014) implemented in the open source software MRtrix(Tournier et al., 2019). Several preprocessing steps utilized FSL(Jenkinson et al., 2012; Smith et al., 2004). Diffusion images were denoised(Veraart et al., 2016), corrected for Gibbs ringing(Kellner et al., 2016), susceptibility distortions(Smith et al., 2004), subject motion(Andersson et al., 2016), and eddy currents(Andersson & Sotiropoulos, 2016). All images were upsampled to of 1.3⨉1.3⨉1.3 mm, and skull-stripping was performed using the Brain Extraction Tool(Jenkinson et al., 2012). Response functions were generated(Dhollander et al., 2016) from both experimental data and the NTU-DSI-122 template at b=1538 s/mm^2^ and b=3077 s/mm^2^ and used to generate FODs(Jeurissen et al., 2014). The number of directions was sufficient to generate FODs with a harmonic order of *l_max_* = 4, which has been suggested to be optimal for registration(Raffelt et al., 2011). Extracellular isotropic CSF-like free water signal fractions were calculated directly from the FODs using 3-tissue constrained spherical deconvolution (3T-CSD), a method that measures cellular microstructure within each voxel fitting into intracellular anisotropic (ICA, WM-like), intracellular isotropic (ICI, GM-like), and extracellular isotropic (ECI, CSF-like/Free Water) compartments(Newman et al., 2020).

Each of the 5 diffusion MRIs were then registered from native space to MNI space using both ANTs SyN algorithm(Avants et al., 2014) to register the processed b=0 s/mm^2^ images with the Colin 27 T2 template(Fonov et al., 2011) and using MRtrix(Tournier et al., 2019) to register the WM FODs with the NTI-DSI-122 template WM FODs(Raffelt et al., 2011, 2012). Each registration was used to generate a transform that was applied to the CSF-like free water signal fraction from that subject. Because these registrations are not deterministic processes, and can produce slightly different transforms each time it is performed on the same images, this procedure was repeated independently 5 times for each subject, via each method (Figure 1).

**Figure 1:**
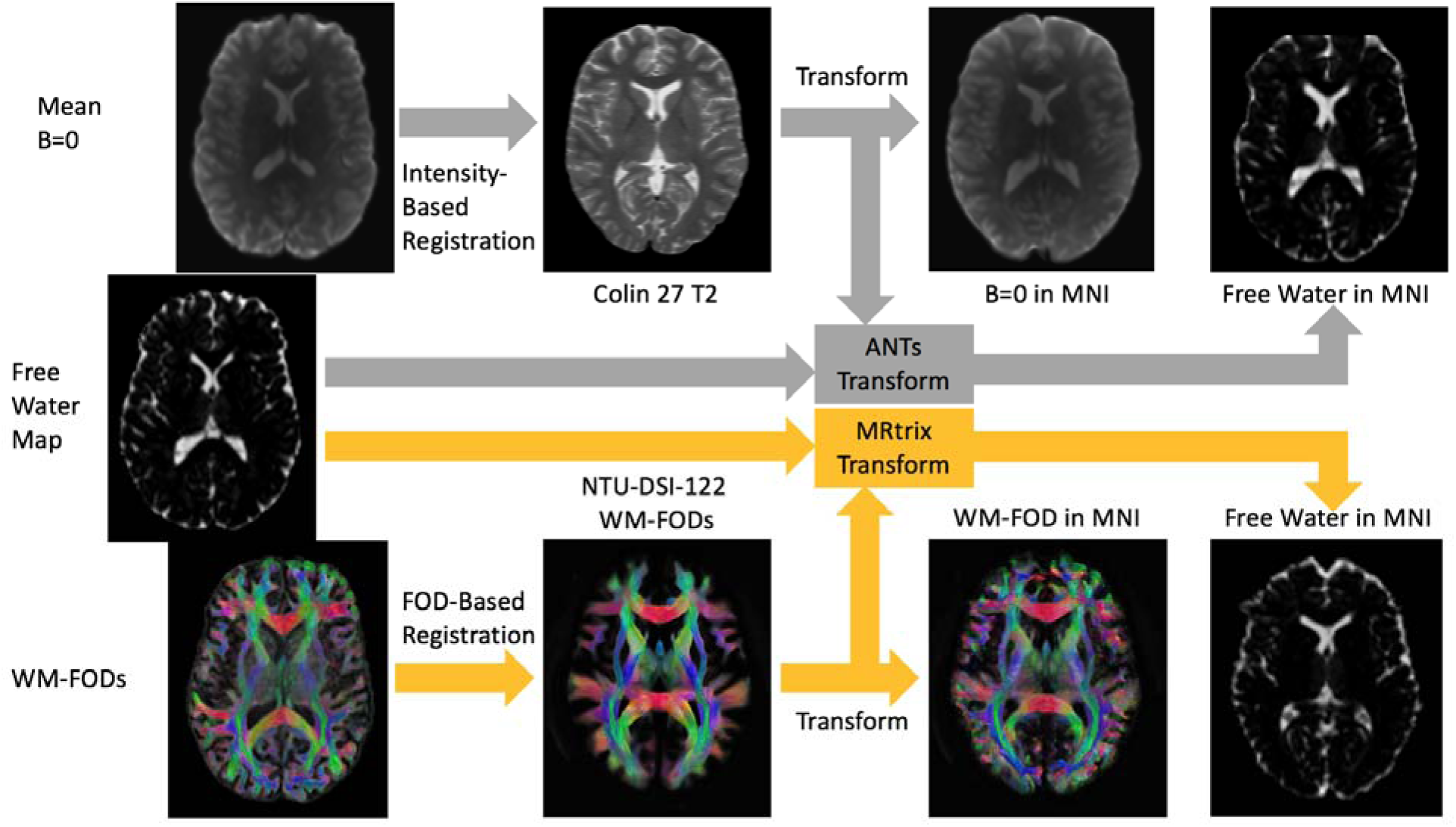
Study workflow illustrating the process of moving from native space images (three images on left), registering with the respective template for each method, applying the transform generated from that registration to the free water signal fraction map, and obtaining the free water signal fraction maps in stereotaxic MNI space (two images on right). This process was repeated 5 independent times for each of the 5 subjects for Experiment 1. In Experiment 2 this process was performed only once for each subject, and the resulting transform was used to move intracellular anisotropic (WM-like), intracellular isotropic (GM-like), and extracellular isotropic (CSF-like/Free Water) signal fraction maps to MNI space.

### Experiment 2: Assessment of Accuracy

To assess the accuracy of each transform method we devised a new method for comparing signal fraction maps in stereotaxic space. A typical approach for registering structural MRIs using b=0 or T1-weighted images has less information for discriminating between voxels within the same tissue type (WM/GM/CSF). FODs however contain a great deal more information in both directional components and axonal fiber density(Raffelt et al., 2011). We hypothesized that this additional information would lead to superior within-tissue registration of cellular microstructure signal fraction maps (as these are derived from FOD coefficients) and thus more accurate whole-brain registration. To test this hypothesis, we generated cellular microstructure signal fraction maps from a ‘ground-truth’ in stereotaxic space by applying the 3T-CSD analysis to the NTU-DSI-122 template itself. Each subject image and the ‘ground-truth’ image could then be thresholded at different signal fraction values and in different directions (the threshold representing either an upper or lower based voxel-wise value limit, illustrated in Figures 2-4). The transformed and thresholded signal fraction image from each subject and from the appropriate tissue type could then be compared for accuracy at multiple levels of detail using the Sorensen-Dice coefficient, a metric of image overlap(Dice, 1945; Sorensen, 1948). By thresholding the subject images after transformation the experiment is also able to account for the impact of interpolation on signal fraction maps. Both MRtrix and ANTs implement a nearest neighbor algorithm, however the greater local deformation that occurs can cause more severe distortions in values. So, while necessary to account for warped voxels, local interpolation changes can be severe for signal fraction maps with value ranges between 0-1. By creating threshold cutoffs between 0.10 or 0.25, large shifts from the ground truth values will cause regions to cross the cutoff threshold and the Dice coefficient value will be lower.

**Figure 2:**
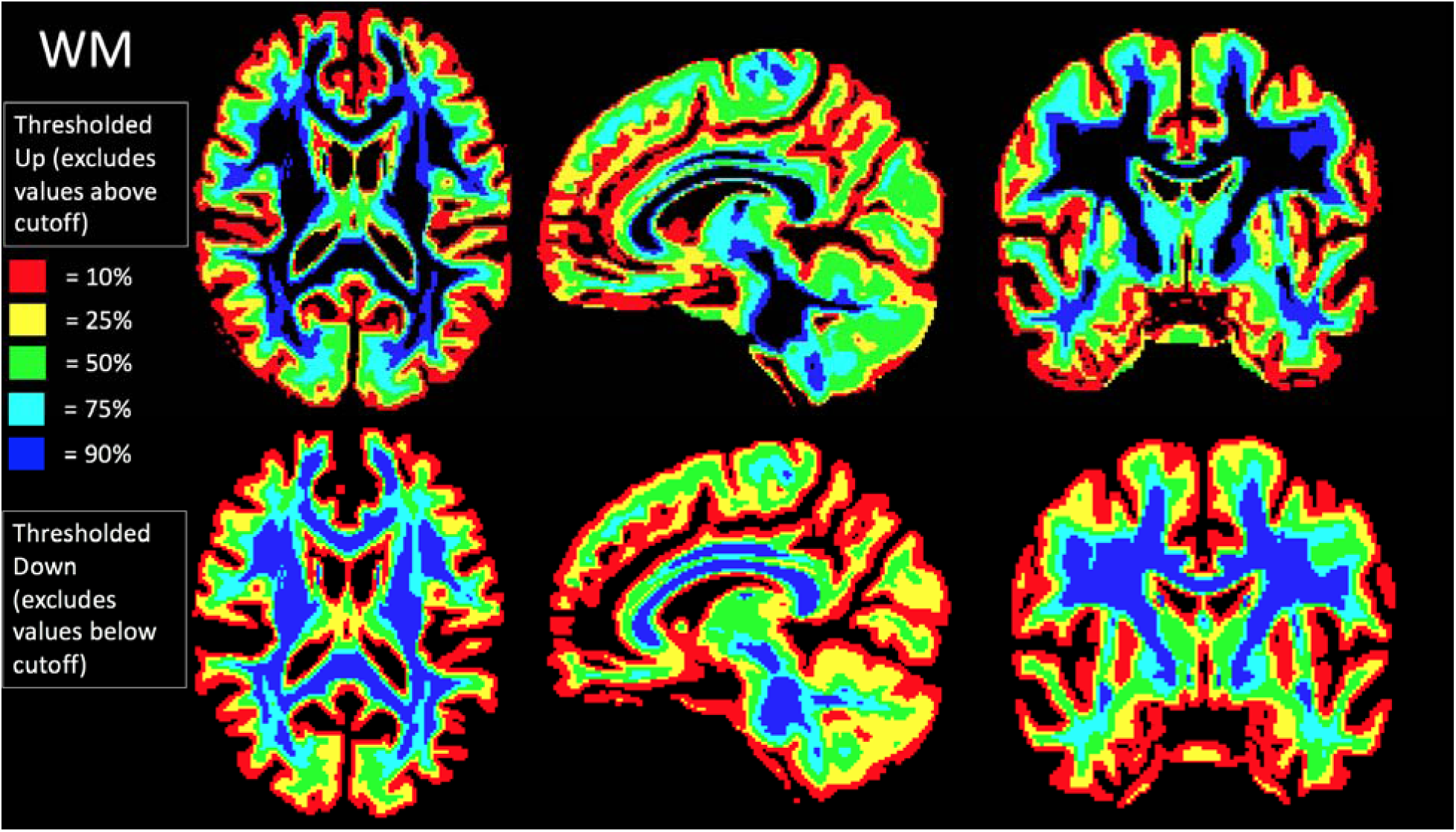
Map of the thresholding procedure for the intracellular, anisotropic (WM) signal fraction map. This is illustrated using the NTU-DSI-122 template which served as a ground-truth structural division in this study. The different direction of thresholding created 10 different ground truth maps for each tissue type, each covering different regions of the brain. For example, the region thresholded ‘Up’ at voxels where the WM signal fraction value was 25% includes both the red and yellow regions largely in the cortex where intracellular anisotropic signal is low, meanwhile the region thresholded ‘Down’ includes the yellow, green, cyan, and blue regions including deep WM but ecluding the outer cortex.

**Figure 3:**
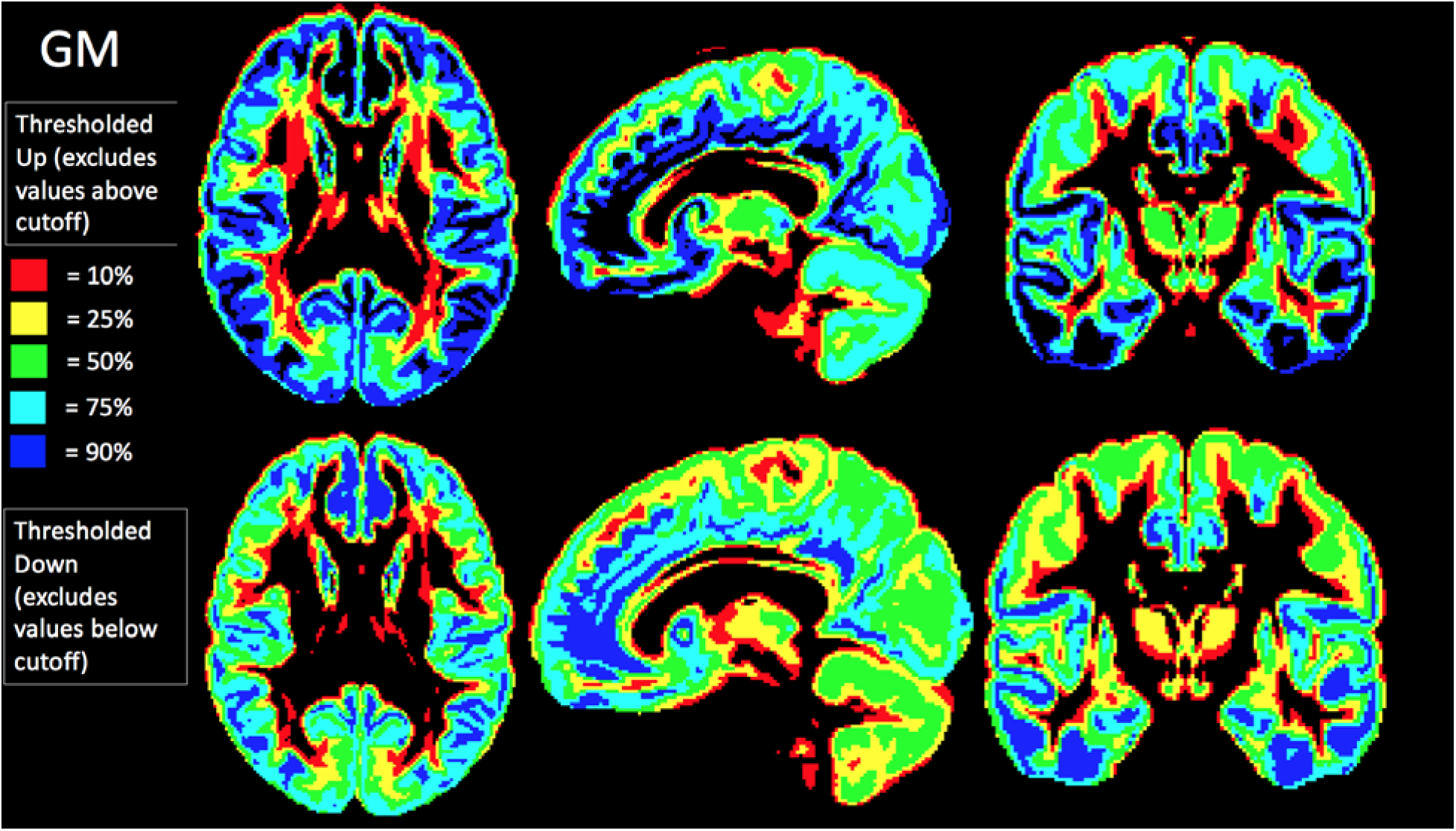
Map of the thresholding procedure for the intracellular, isotropic (GM) signal fraction map. This is illustrated using the NTU-DSI-122 template which served as a ground-truth structural division in this study. The different direction of thresholding created 10 different ground truth maps for each tissue type, each covering different regions of the brain. This patterning includes portions of central structures as well as detailed layering of the cortex without a large contribution from axonal WM areas.

**Figure 4:**
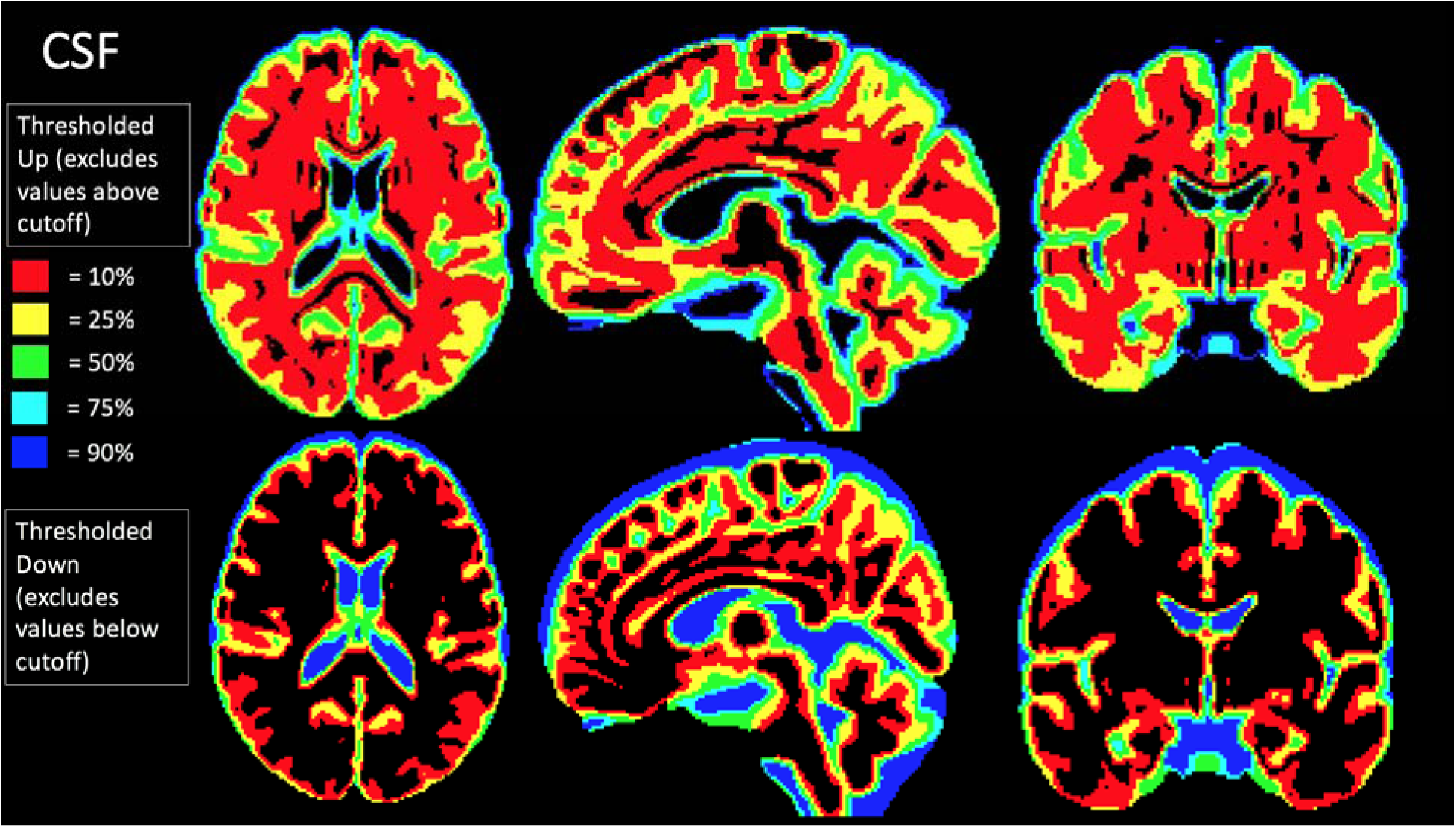
Map of the thresholding procedure for the extracellular, isotropic (CSF/free water) signal fraction map. This is illustrated using the NTU-DSI-122 template which served as a ground-truth structural division in this study. The different direction of thresholding created 10 different ground truth maps for each tissue type, each covering different regions of the brain. This patterning includes detailed ventricles as well as a highly detailed layering of the cortex displaying the gradient of fluid infiltration into cortex toward the outer edge of the brain parenchyma.

For subjects, diffusion MRI images were obtained from a random selection of 100 non-twin individuals in the Human Connectome Project (HCP)(Van Essen et al., 2013). Subjects were scanned using a specially modified Siemens Skyra 3T scanner with acquisition parameters of 1.25×1.25×1.25mm^3^ isotropic voxels, TE=89ms and TR=5520ms; 36 b=0 images were acquired interleaved with 180 gradient directions each at b=1000s/mm^2^, b=2000s/mm^2^, and b=3000s/mm^2^(Sotiropoulos et al., 2013; Van Essen et al., 2013). Processing of dMRI data was performed identically to Experiment 1 with the exception of single-shell constrained spherical deconvolution following preprocessing(Dhollander & Connelly, 2016). Quality control was performed via manual visual inspection of each subject’s signal fraction maps to remove subjects who experienced registration failure (largely blank final images or the presence of extreme distortions) and 4 subjects were removed from the study (three for failure of the FOD-based method and one for failure of the ANTs method).

The closest matching non-b=0 shell was extracted and FODs were calculated as before. Using the NTU-DSI-122 allowed for the creation of a ‘ground-truth’ signal fraction map for each of the tissue types derived from 3T-CSD (ICA, ICI, and ECI). For the Colin-27 template used as an example comparison tissue segmentation was performed using FSL(Jenkinson et al., 2012).

No additional preprocessing was necessary and response functions were derived from the dMRI image itself following the protocol established in Experiment 1.

### Statistical Approach

In Experiment 1 once the free water signal fraction maps had been moved into stereotaxic space the mean squared difference was then calculated for each whole-brain image between every same-method same-subject combination to analyze which method more consistently transformed the free water signal fraction map into MNI space. This approach allowed for transform reliability to be assessed using traditional statistical approaches.

In Experiment 2 Sorensen-Dice coefficients were calculated for each of the subjects’ signal fraction maps that passed quality control. Each subject had 5 cutoff maps per direction of thresholding (upper thresholding above the cutoff value and lower thresholding below the cutoff value) for each of the three tissue types derived from 3T-CSD (ICA, ICI, and ECI) in addition to whole brain maps (compared against both NTU-DSI-122 and Colin27 ground truth signal fraction or determinative tissue divisions) for a total of 36 measurements per subject. Each direction of thresholding and each tissue type was compared between registration methods to determine if registration using intensity or FOD-based methods were consistently more successful at correctly aligning each level of the signal fraction maps.

## Results

### Experiment 1: Assessment of Reliability

Each of the subject’s calculated mean squared difference was lower for the MRtrix FOD transform compared to the ANTs SyN tansform. This resulted in a significantly lower mean squared difference between repetitions compared to the ANTs SyN generated transform (pairwise T-test, T_49_ = 8.02, p<0.001) indicating that the FOD algorithm was able to more consistently register and transform the free water signal fraction maps (Figure 5).

**Figure 5:**
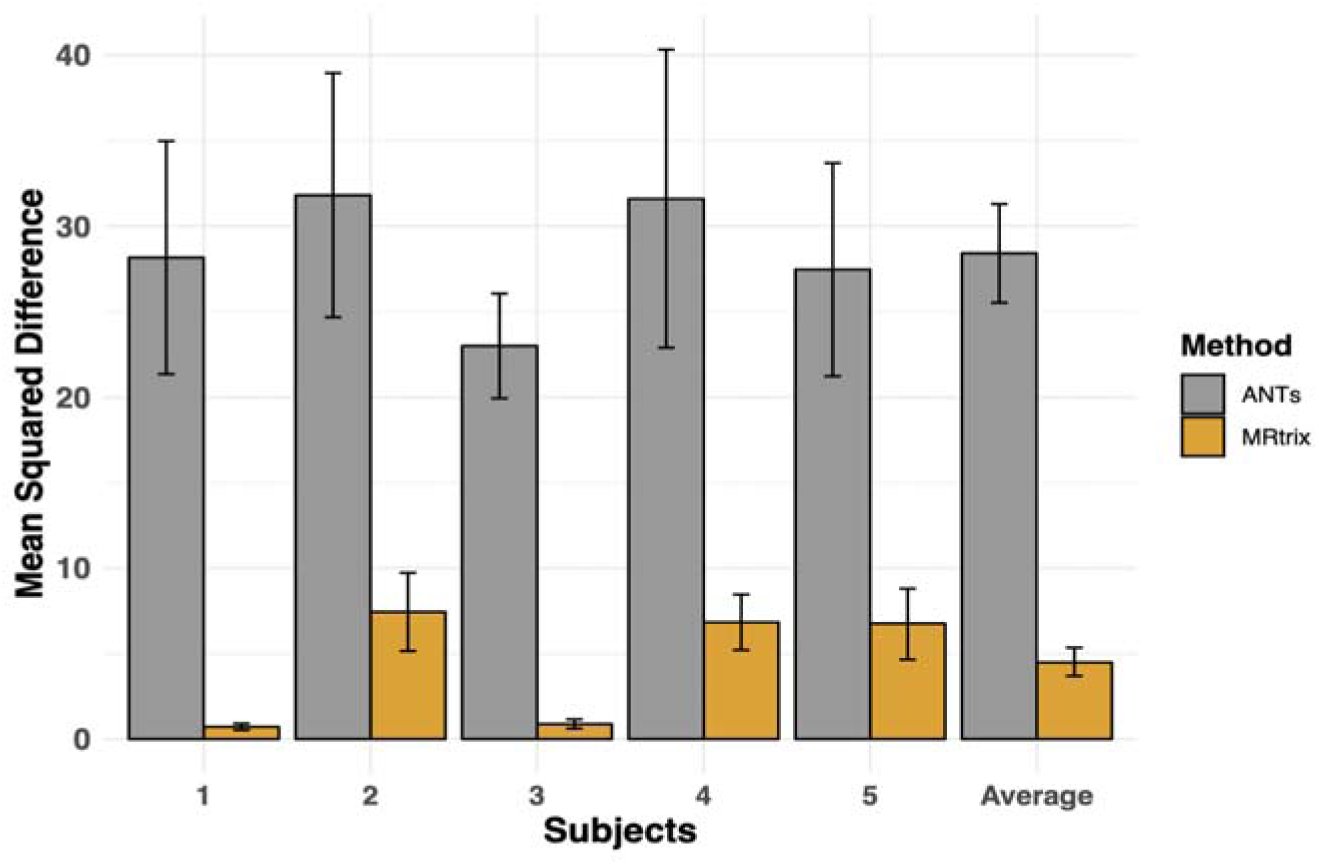
The mean squared difference results from Experiment 1 between each of the 5 independent registration attempts for each of the 5 subjects involved in analysis are presented, alongside the group mean (±SE). Grey bars represent the SyN intensity-based registration implemented in ANTs and yellow bars represent the WM-FOD based registration implemented in MRtrix. The reliability of the MRtrix implemented FOD-based registration method had a significantly lower mean squared difference between the transformed images of each subject (pairwise T-test, T_49_ = 8.02, p<0.001).

### Experiment 2: Assessment of Accuracy

Registration using the FOD-based template resulted in signal fraction maps with a greater Sorenson-Dice coefficient in upper thresholded ICA (Repeated measures ANOVA, F_1,950_=28485.01; p<0.001), upper thresholded ICI (Repeated measures ANOVA, F_1,950_=26033.74; p<0.001), upper thresholded ECI (Repeated measures ANOVA, F_1,950_=11189.9; p<0.001), lower thresholded ICA (Repeated measures ANOVA, F_1,950_=3600; p<0.001), and lower thresholded ICI (Repeated measures ANOVA, F_1,950_=23.5, p<0.001). Intensity-based registration implemented using ANTs resulted in signal fraction maps with a greater Sorenson-Dice coefficient in lower thresholded ECI (Repeated measures ANOVA, F_1,950_=892.8; p<0.001). When each directional threshold was combined FOD-based registration had a significantly higher Dice coefficient for each tissue type (ECI Repeated measures ANOVA, F_1,1910_=25.78; p<0.001; ICI Repeated measures ANOVA, F_1,1910_=35.36; p<0.001; ICA Repeated measures ANOVA, F_1,1910_=40.26; p<0.001) These results are summarized in Figure 6. Whole brain registration results were compared by calculating Dice coefficients for both the 3T-CSD derived tissue signal fraction maps and the FSL intensity based determinative tissue segmentations.

**Figure 6:**
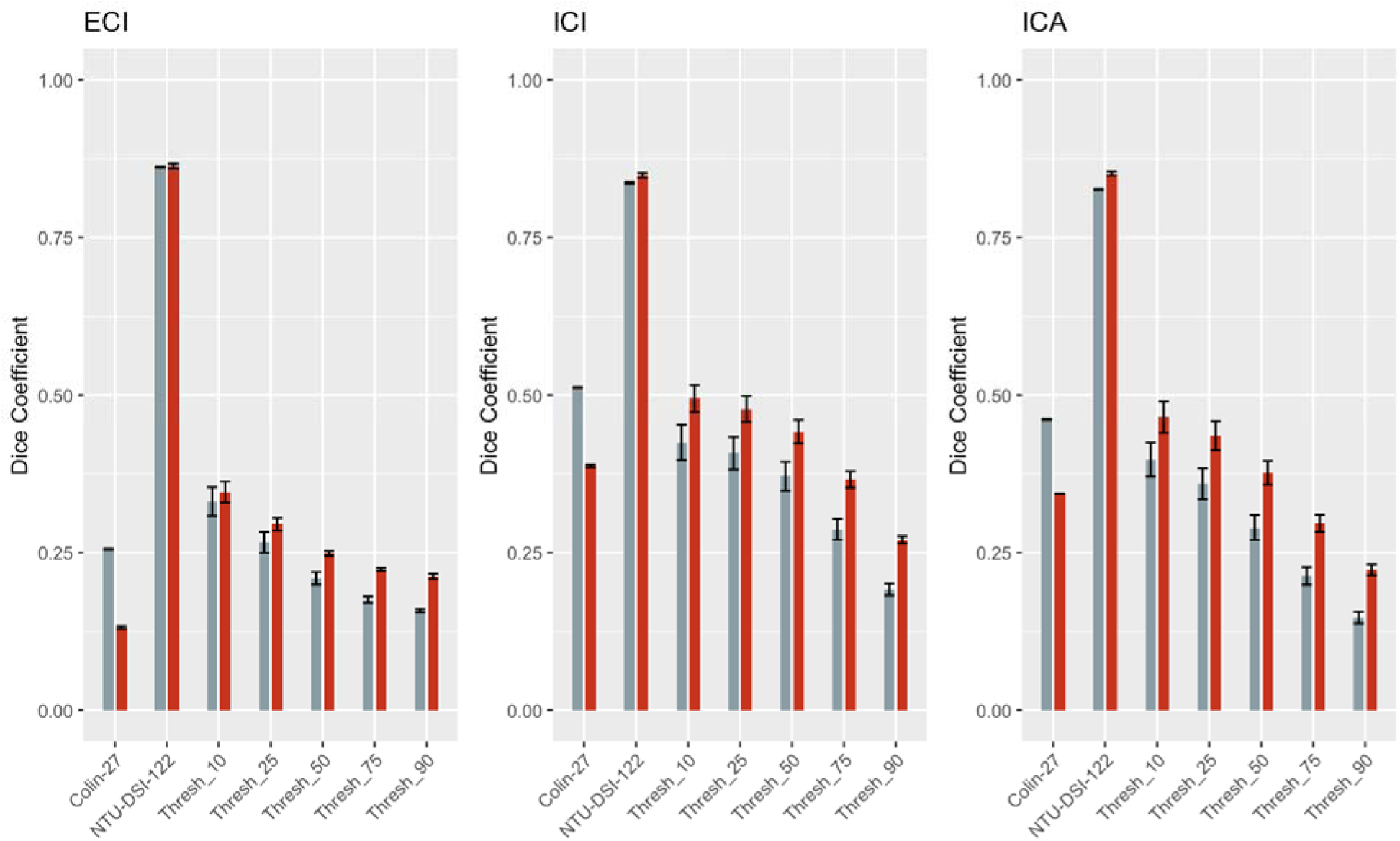
Chart displaying the Dice coefficient calculated for each tissue type and averaged across all subjects and threshold levels, and compared to the ground truth of either the Colin-27 binary tissue division (Gray; CSF/GM/WM) or the NTU-DSI-122 signal fractions (Red; ECI/ICI/ICA). Dice coefficient calculated from the intensity-based registration method is displayed in gray (±SE) and the FOD-based registration method is displayed in red (±SE).

There was a significant effect of registration method on Dice coefficient (Fig. 6), with FOD-based registration performing better than intensity-based registration on each NTU-DSI-122 comparison while intensity-based registration had a higher dice coefficient when compared to the Colin-27 segmentations, however the NTU-DSI-122 comparison was far higher for both methods (Repeated measures ANOVA, F_1,1146_=12.69; p<0.001). There was also a significant effect of tissue type (Repeated measures ANOVA, F_1,1146_=21.77; p<0.001), with the ECI free water tissue compartment having the lowest Dice coefficient for both methods when compared to the Colin-27 segmentation and the highest Dice coefficient for both methods when compared to the NTU-DSI-122 template. There was no significant interaction between registration method and tissue type (Repeated measures ANOVA, F_1,1146_=0.085; p=0.918 n.s.).

## Discussion

We have demonstrated the successful use of an FOD-based template for the reliable and accurate transformation of signal fraction maps into stereotaxic space. Using the NTU-DSI-122 we are able to create an FOD template suitable for individual or template-based registration. The NTU-DSI-122 is a flexible template with a variety of b-value shells to create suitable b-value matched templates to the majority of common acquisition schemes. The additional directional information inherent in the FOD map provides a powerful means to register axonal fibers and to generate within-tissue contrast that is not possible using regular structural imaging techniques.

We have compared this FOD-based template registration technique to the widely used intensity-based SyN transform implemented in ANTs(Avants et al., 2014; Klein et al., 2009). Registering dMRI images using the NTU-DSI-122 template with an FOD-based apodized point spread function(Raffelt et al., 2012) was demonstrated to be more reliable and, in a variety of contexts, more accurate at registering brain microstructure maps than the intensity-based method. The NTU-DSI-122 showed better whole brain registration and accuracy at various thresholds and in each of the tissue microstructure types examined. This is especially notable due to only information relating to the anisotropic tissue compartment (ICA/WM) being used in the registration process. Intensity-based registration uses information from each of the different tissue compartments, and so it would not have been unexpected to have performed better at the GM/CSF boundary at the edge of the cortex however each of the tissue compartments had a higher Dice coefficient when FOD-based registration was used. This finding supports the view that each tissue compartment contributes some signal to almost every voxel, in contrast to a binary view where voxels belong to a specific predominant tissue type. Perhaps unsurprisingly, the ECI compartment performed the worst in regards to accuracy at various thresholded levels between both FOD- and intensity-based registration methods. Interestingly however, the FOD-based registration outperformed the intensity-based registration method throughout the mid-range of thresholded values which suggests that the FOD-based method was more accurate at registering the sulci and gyri (the largest contributors to these thresholds as illustrated in Figs. 2-4).

The increased accuracy of the FOD-based registration method is complemented by the range of b-value shells present in the NTU-DSI-122 template, allowing for a template to be created that matches experimentally acquired acquisitions. In this study we have exclusively tested registering dMRI images from natively acquired subject space to stereotaxic space, while this is not necessarily the optimal method for every study, it allowed for an easy to analyze experimental design and presented a reasonably challenging task for a registration algorithm to align subject brains with individual variations to a normalized template. A common alternative to direct native space to stereotaxic space registration is to create a cohort specific template to first register all subjects to, then performing a single registration between that template and stereotaxic space(Newman et al., 2020; Raffelt et al., 2011). To illustrate the utility of the NTU-DSI-122 and the suitability of this method to register cohort specific templates from a variety of acquisitions a number of templates from a wide variety of subject cohorts and acquisitions have been warped into stereotaxic space. The FODs from these templates in MNI space are illustrated in Figure 7, they display accurate anatomical registration from cohorts across the lifespan and including clinical quality data.

**Figure 7:**
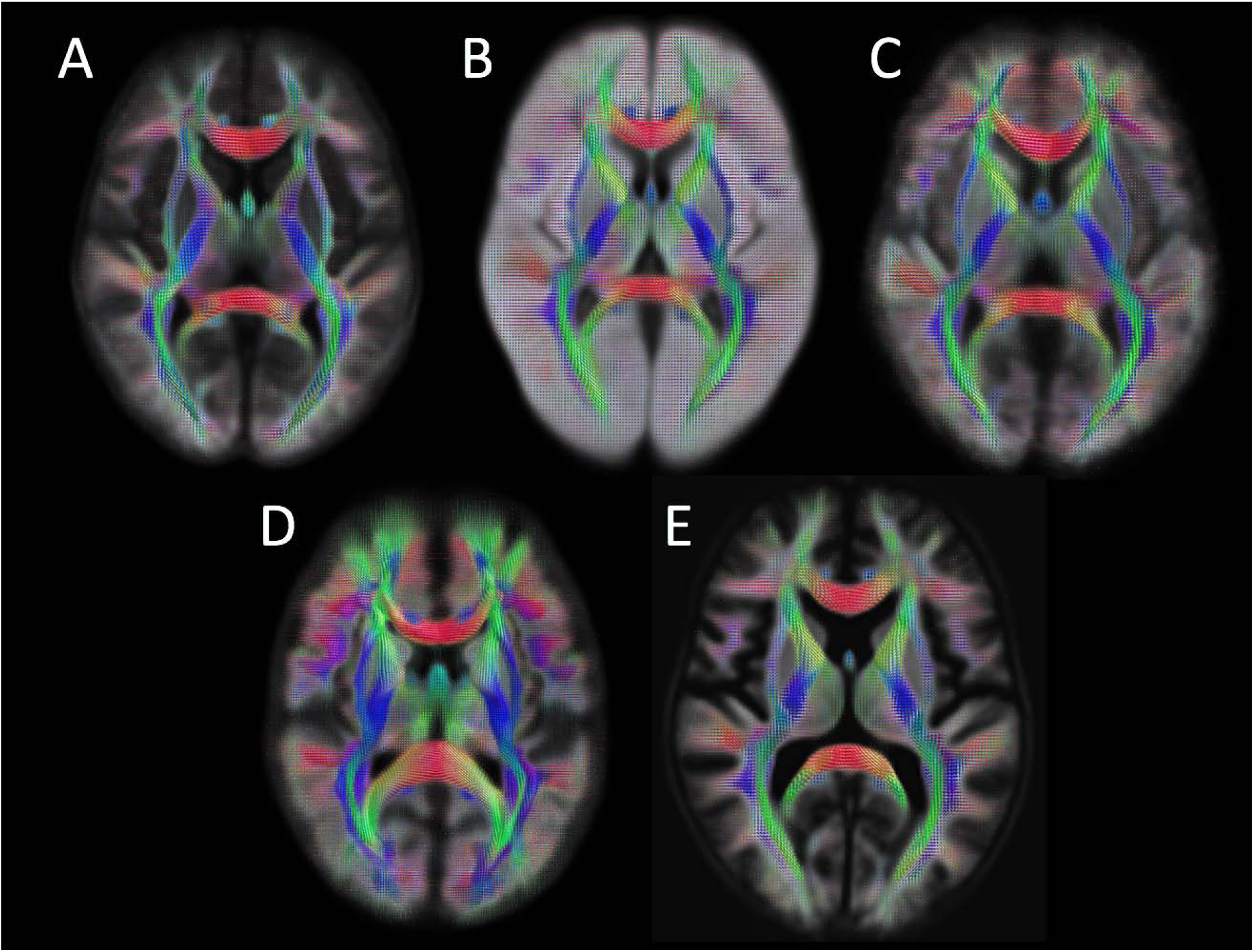
Illustration of 5 cohort-specific templates transformed into MNI space using b-value matched variants of the NTU-DSI-122. Each template was constructed from between 30-50 randomly selected individuals from their respective studies. Each cohort was collected with a different acquisition protocol (both single- and multi-shell acquisitions ranging from b=1000 s/mm^2^ to b=3000 s/mm^2^ maximum b-value) across different age groups and template B was constructed from a clinically acquired dataset. Cohort age ranged from 9-11 years old (A), 41-84 years old (B), 32-37 years old (C), 18-27 years old (D), and 56-82 years old (E).

The purpose of this study is to introduce a new template option for FOD-based registration of dMRI microstructure measurements to stereotaxic space. To that end, this study as much as possible strived to maintain a simple and straightforward registration pipeline that would mimic a typical user’s ‘out-of-the-box’ experience using the neuroimaging software available with MRtrix and ANTs(Avants et al., 2014; Tournier et al., 2019). There are a number of ways to increase registration algorithm performance by altering the settings of these algorithms and optimizing the algorithm to work with data from a particular pipeline or even from a specific subject. For example, the accuracy of both registration methods tested here can sometimes be improved if the number of iterations allowed before the algorithm terminates is increased, or if the hierarchical resolution steps are modified. While these are important considerations for implementing registration in any specific study, we aimed to demonstrate a simple comparison for validity, not to specify the total superiority of one method over another. Thus, optimizing the algorithms is beyond the scope of this paper.

Intensity-based registration has a number of advantages over an FOD-based method that were not evaluated. This study utilized two datasets with high-quality dMRI acquisitions intended for research applications, one collected locally and another from the HCP(Sotiropoulos et al., 2013; Van Essen et al., 2013). The high b-value shells and number of gradient directions are well-suited for generating FOD maps with sufficiently high ϑmax spherical harmonic degree to detect multiple directions of crossing WM fibers(Genc et al., 2020; Jeurissen et al., 2014). A clinically acquired dataset or dMRI with only a limited number of directions intended for diffusion tensor imaging (DTI) may not be suitable for FOD-based registration or may suffer decreased accuracy while intensity-based registration only requires a single b=0 image. This study also does not test the ability of intensity-based registration techniques to register single volume DTI derivative metrics, such as fractional anisotropy, to a template or stereotaxic space.

Given the rise in cellular microstructure models of dMRI data, it is imperative that well evaluated registration methods are used to ensure accurate and reliable transforms. This will only become more important as large cohort datasets require automated processing pipelines with minimal user intervention. The addition of multi-site data provides another imperative to have reliable registration that does not introduce a new source of variation into the study. This study describes a novel registration process that uses the NTU-DSI-122 as a FOD template located in stereotaxic space suitable for FOD-based registration. This template and the software used in this study are freely available and available for use by any dMRI researcher.

### Conclusion

Using the NTU-DSI-122 as a template for FOD-based registration provides a means to register subject brain scans to stereotaxic space. This method is more reliable and more accurate than a leading intensity-based registration method at registering maps of brain cellular microstructure.

